# Systemic benefits of Gc inhibition to preserve insulin sensitivity

**DOI:** 10.1101/2020.10.06.328963

**Authors:** Taiyi Kuo, Domenico Accili

## Abstract

Type 2 diabetes is caused by an imbalanced supply and demand of insulin. Insulin resistance and impaired β-cell function contribute to the onset of hyperglycemia. No single treatment modality can affect both aspects of diabetes pathophysiology. Thus, current treatments focus either on increasing insulin secretion (incretin mimetics, sulfonylureas) or insulin sensitivity (metformin and TZD), or reducing hyperglycemia (insulin, sglt2i). Previously, we reported that ablation of *Gc*, encoding a secreted protein with a primary role in vitamin D transport, improves pancreatic β-cell function in models of diet-induced insulin resistance. Here, we show that *Gc* ablation has systemic insulin-sensitizing effects to prevent weight gain, hyperglycemia, glucose intolerance, and lower NEFA and triglyceride in mice fed a high-fat diet. Hyperinsulinemic-euglycemic clamps show that Gc ablation protects insulin’s ability to reduce hepatic glucose production, and increases glucose uptake in skeletal muscle and adipose tissue. Moreover, acute Gc inhibition by way of adeno-associated virus encoding a short hairpin RNA to promote Gc mRNA degradation, prevents glucose intolerance caused by high fat feeding. The data suggest that Gc inhibition can provide an approach to increase insulin production in β-cells, and insulin action in peripheral tissues.

**RESEARCH IN CONTEXT:**

▪ The goal was to find a therapeutic target that can improve insulin sensitivity and β-cell function simultaneously.
▪ Gc ablation preserves β-cell insulin secretion *ex vivo* and *in vivo*.
▪ Deletion of Gc prevents weight gain, reduces fat mass, lowers fasting glycemia, improves glucose tolerance, reduces hepatic glucose production after feeding, and increased glucose uptake in muscle and adipose.
▪ Acute Gc inhibition improves glucose tolerance, which suggests that targeting Gc could provide an alternative way to treat type 2 diabetes.

## INTRODUCTION

Insulin resistance and β-cell failure underlie most forms of type 2 diabetes, and efforts have been directed to increase insulin production and insulin sensitivity as diabetes treatments (1). During the natural history of type 2 diabetes, insulin resistance possibly antedates β-cell abnormalities but remains relatively constant (2,3), whereas β-cell dysfunction deteriorates rapidly following the onset of hyperglycemia (4,5). Advances in understanding the pathophysiology have shaped diabetes treatment: combinations of agents that improve β-cell function with those that decrease insulin resistance are now considered standard therapy (6). In recent years, incretin-based approaches have all but replaced sulfonylureas as insulin secretagogues (7), with additional benefits on a variety of tissues and cell types (8). On the other hand, the development of effective insulin sensitizers has lagged behind, despite strong evidence suggesting that they are effective at preventing diabetes (9-11).

*Gc* (encoding Group-specific component, or vitamin D binding protein) is expressed primarily in liver and is secreted into the bloodstream, where its primary function is to act as a cargo for vitamin D and its metabolites (12). Interestingly, its expression in the pancreas is limited to glucagon-producing α-cells (13). However we discovered that, during the progression of multiparity- or age-induced diabetes in mice, Gc expression is ectopically activated β-cells (14). Moreover, in the absence of β-cell transcription factor Pax6 or Isl1, Gc expression was elevated in β-cells or islets, respectively (15,16). While its function in the endocrine islet remains unclear, genetic knockout of Gc preserved insulin secretory response and β-cell function *in vivo*, as determined by hyperglycemic clamps, and *ex vivo* in isolated islets (14).

In the course of experiments using high-fat diet (HFD) to induce insulin resistance and hyperglycemia and investigate Gc function in the endocrine pancreatic islet, we observed that Gc-deficient mice gained less weight than their control WT littermates. This observation prompted us to conduct an in-depth investigation of the metabolic effects of genetic ablation of Gc on systemic insulin sensitivity and insulin action. Here, we report that Gc ablation preserves insulin sensitivity in liver, skeletal muscle, and adipose tissue in diet-induced obesity, and results in lower weight and fat mass, as well as lower NEFA and triglycerides. In addition, acute Gc inhibition in liver improves glucose tolerance in HFD-fed mice.

## METHODS

### Animal care and use

We generated Gc knockout mice (GcKO) using frozen sperm from the UCD Knockout Mouse Project Repository (Gc^tm1.1(KOMP)Vlcg^) (14). Mice were weaned at 3 weeks of age, housed at 22-24°C, fed normal chow diet (NCD) or high fat diet (HFD), and maintained on a 12-hr light-dark cycle (lights on at 7 AM). Based on caloric content, NCD consists of 13% fat, 62% carbohydrates, and 25% protein (PicoLab rodent diet 20, 5053; Purina Mills), while HFD contains 60% fat, 24% carbohydrates, and 16% protein (Research Diets, D12492). Littermate control mice for GcKO retained at least 1 WT *Gc* allele. The Columbia University Institutional Animal Care and Utilization Committee approved all animal procedures.

### Metabolic analyses and indirect calorimetry

We performed intraperitoneal glucose tolerance tests (ipGTT) by injecting glucose (2g/kg for NCD-fed mice, or 1g/kg for HFD fed mice) after an overnight fast. Insulin tolerance tests were performed by injecting insulin (0.75 units/kg) after a 4-hr fast, and intraperitoneal pyruvate tolerance tests by injecting sodium pyruvate (2g/kg) after an overnight fast (17). We measured insulin levels with an ELISA kit (Mercodia) and glucagon levels with radioimmunoassay kit (MilliporeSigma). We estimated body composition by nuclear magnetic resonance (Bruker Optics). We measured food intake, energy expenditure, and respiratory exchange rate (RER) with a Comprehensive Laboratory Animal Monitoring System (CLAMS, Oxymax) (Columbus Instruments, OH, USA). Plasma triglyceride (TG), total cholesterol E and non-esterified fatty acid (NEFA) were measured with commercially available kits following manufacturers’ instructions: Infinity TG kit (Thermo Fisher Scientific), total cholesterol E kit (Wako Diagnostics), and NEFA kit (Wako Diagnostics).

### RNA preparation and quantitative PCR

We isolated total RNA with Nucleospin RNA kit (Macherey-Nagel), and followed previously described protocol for reverse transcription (18). We used the resulting cDNA to perform quantitative PCR with GoTaq master mix (Promega), and analyzed data with the standard ΔΔCt method. Rpl19 was used for internal normalization. Primer sequences are listed in Supplemental Table S1.

### Western blotting and imaging

We used previously described protocol for immunoblotting (19). A 30-mg liver sample was placed in 600 μl RIPA lysis buffer (Millipore, 20-188), supplemented with Halt protease and phosphatase inhibitor cocktail (Thermo Scientific, 78440). The sample was homogenized for 30 seconds, on setting 6 of an IKA T10 Basic Ultra-Turrax homogenizer system. Then, samples were spun down using a benchtop centrifuge at 4°C with 20,817 rcf for 10 min, and supernatant was transferred to a new tube avoiding the lipid layer and precipitant. The supernatant was sonicated for a total of 5 min (10 sec on and 10 sec off for 10 min) on setting 9 of a model 550 Sonic Dismembrator (Fisher Scientific). The supernatant was spun again and transferred to a new tube avoiding precipitant, followed by protein concentration measurement (Pierce BCA protein assay kit).

The following primary antibodies were used: anti-Gc (HPA019855, Atlas Antibodies, 1:1000 dilution); anti-total Akt (Cell Signaling, no. 9272, 1:1000 dilution); anti-phosphorylated Akt at serine 473 (Cell Signaling, no. 4060, 1:500 dilution); and anti-β actin (abcam, ab8227, 1:1000 dilution). IRDye 800CW (LI-COR, no. 925-32211) or IRDye 680LT (LI-COR, no. 925-68021) goat anti-rabbit secondary antibody was used. Proteins were detected using an Odyssey imaging system (LI-COR), and quantifications were measured using ImageJ.

### Hyperinsulinemic-euglycemic clamps

We performed hyperinsulinemic-euglycemic clamps as previously described (20). Briefly, during the 120-min clamp period, insulin was infused at a constant rate of 2.5 mU/kg/min to raise circulating insulin levels about 3-fold over basal levels. Concurrently, glucose was infused intravenously at variable rates to maintain euglycemia, and this rate of glucose infusion directly correlates with insulin sensitivity. Plasma glucose concentrations were measured every 10 to 20 min from the tail vein to adjust the rates of glucose infusion needed to maintain a constant glycemia. Furthermore, radioactively labeled [3-3 H] glucose (18 μCi per mouse) and 2-deoxy-D-[1-14 C] glucose (a bolus of 10 μCi per mouse) were used to measure hepatic insulin action and glucose metabolism in skeletal muscle and adipose tissues, respectively.

Prior to the 120-min clamp period, [3-3 H] glucose (0.05 μCi/min) was infused for 120 min to examine the basal rate of whole-body glucose turnover. Plasma samples were taken at the end to measure basal levels of insulin, plasma glucose, [3H] glucose, specific activity, and rates of glucose turnover. During this basal state, hepatic glucose production is the only source where the glucose can be introduced into the circulation, therefore represents a steady state for peripheral glucose disposal. As a result, the basal rate of whole-body glucose turnover equilibrates with the basal rate of hepatic glucose production.

During the 120-min clamp period, [3-3 H] glucose (0.1 μCi/min) continued to be infused to measure insulin-stimulated whole-body glucose turnover rate. Plasma [3H] glucose concentrations recorded during the final 30 min of the clamp were applied to calculate the specific activity and glucose turnover rates. Furthermore, a one-time, non-metabolizable 2-deoxy-D-[1-14 C] glucose infusion at 75 min into the clamp period was used to measure glucose uptake in gastrocnemius muscle and epidydimal white adipose tissue.

Whole-body glycolysis was obtained from the rate of elevated plasma ^3^H2O as a byproduct of glycolysis, calculated with linear regression based on measurement from time point 80 to 120 min with a 5-min interval. Whole body glycogen synthesis was estimated by subtracting whole-body glycolysis from whole-body glucose turnover. At the end of the clamp period, mice were anesthetized using sodium pentobarbital, and tissues were harvested for biochemical analysis.

### Adeno-associated virus injection

Because of liver tropism of adeno-associated virus subtype 8 (AAV8), AAV8-H1-shRNA was chosen as a vector to mediate the knockdown of hepatic Gc (21). We generated AAV8-H1-shRNA against Gc (sh-Gc) to silence hepatocyte Gc, as well as a scrambled shRNA control (sh-scr). Three-month-old WT mice were fed a high fat diet for 12 weeks prior to viral injection. Viral injection was performed through the tail vein at a dose of 1.5 ×10^11^ genome copies per mouse. Glucose tolerance tests were performed prior to, and three weeks post-viral injection.

### Statistical Analysis

Data were analyzed with Prism 8 (GraphPad). Results are shown as mean values ±SEM. *P* values ≤0.05 are considered statistically significant. Statistical analyses were carried out by Student’s *t* test when two groups were compared, and by ANOVA when more than two groups were analyzed.

### Data sharing and availability

The datasets generated and/or analyzed during the current study are available from the corresponding author upon reasonable request.

## RESULTS

### Reduced weight and fat mass in HFD-fed GcKO mice

We have previously shown that Gc ablation benefits β-cell function under conditions of impaired metabolism, such as multiparity or high-fat diet (HFD). This effect reflects a local, possibly paracrine action of Gc in the islet during the progression of diabetes, as its expression, normally restricted to α-cells, becomes activated in β-cells (14). However, the metabolic role of liver-derived plasma Gc, by far its largest fraction, remained unclear. To address this question, we studied the metabolic features of HFD-fed GcKO mice and their WT littermates. Following 12 weeks on HFD, GcKO mice showed ∼19% lower body weight compared to WT (Fig. 1A). We utilized dual-energy X-ray absorptiometry to determine body composition. We found that the lower weight of GcKO mice could be accounted for by a 30% decrease of fat mass (25.6±1.0 *vs*. 17.7±2.9 g, *p* <0.05) (Fig. 1B), while lean mass was unchanged (Fig. 1C). To assess the contribution of food intake to the reduced body weight in GcKO mice, we measured food consumption but found no difference between WT and GcKO mice (Fig. 1D). To determine whether Gc ablation affected energy homeostasis, we subjected mice to indirect calorimetry measurements in metabolic cages, but did not find a significant difference in energy expenditure (Fig. S1A, S1B) or respiratory exchange rate (Fig. S1C-G). However, GcKO mice showed ∼22% decrease of fasting glycemia compared to WT (138.1±5.4 *vs*. 108.1±2.7 mg/dl, *p* <0.001 (Fig. 1E).

**Figure 1.**
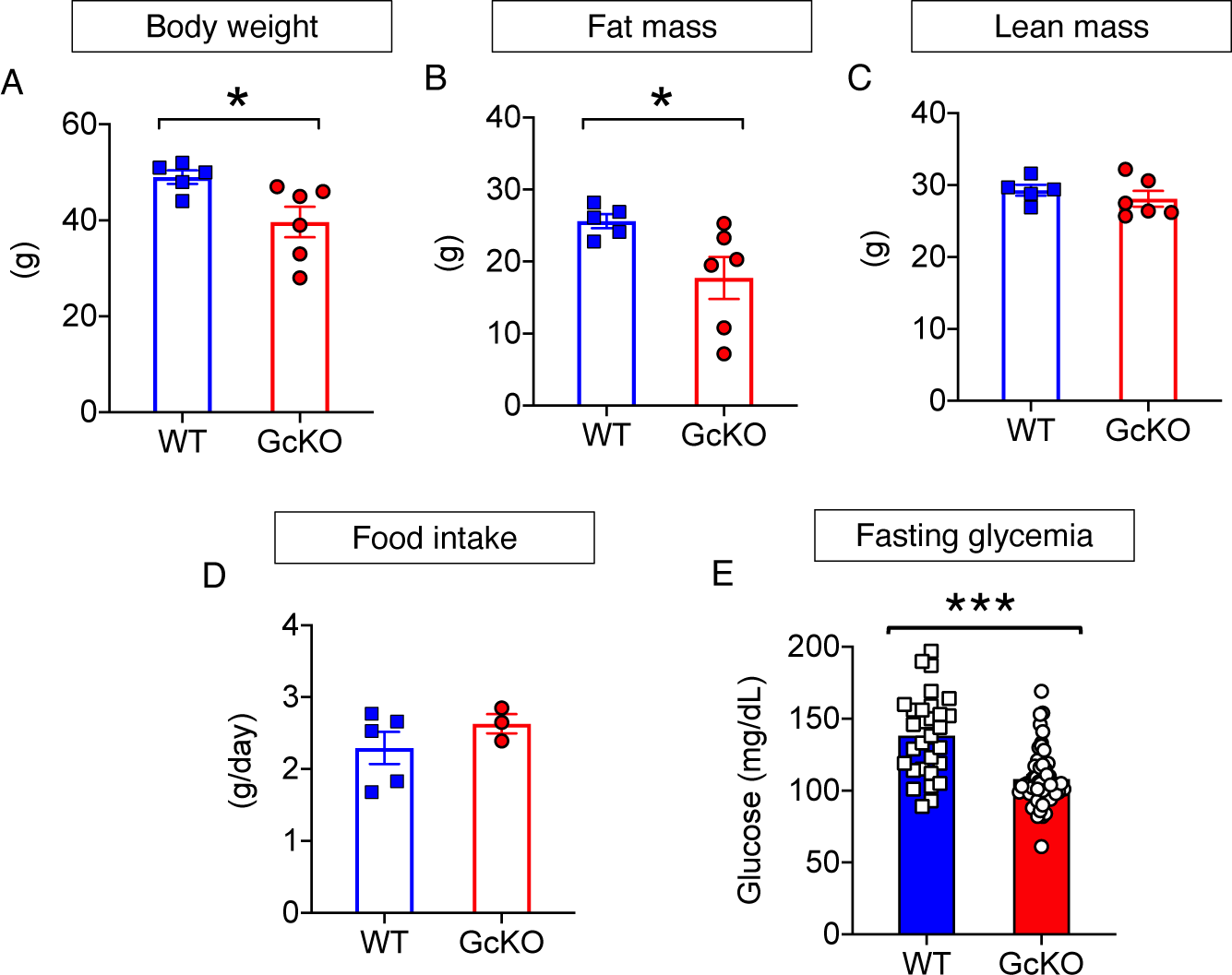
Metabolic features and body composition of HFD-fed mice. **(A)** Body weight, **(B)** fat mass, and **(C)** lean mass of HFD-fed WT and GcKO mice. **(D)** Food consumption of WT and GcKO mice. WT n=5, GcKO n=6 **(E)** Fasting glucose levels of WT and GcKO mice. WT n=29, GcKO n=53. Error bars represent ±SEM, \**p* <0.05, \*\*\**p* <0.005 by Student’s *t* test.

### Deletion of Gc improves glucose tolerance

To evaluate the consequences of Gc ablation on glucose metabolism, we performed glucose tolerance tests in chow-or HFD-fed GcKO and WT mice. HFD resulted in a significant deterioration of glucose tolerance in male, but not in female WT mice (Fig. 2A-B and S2A-B). We observed a significant improvement of glucose tolerance in HFD-fed male GcKO mice compared to WT (Fig. 2A-B). In contrast, no difference was observed in mice on chow diet (Fig. S2C-F). Therefore, we focused further experiments on HFD-fed male mice.

**Figure 2.**
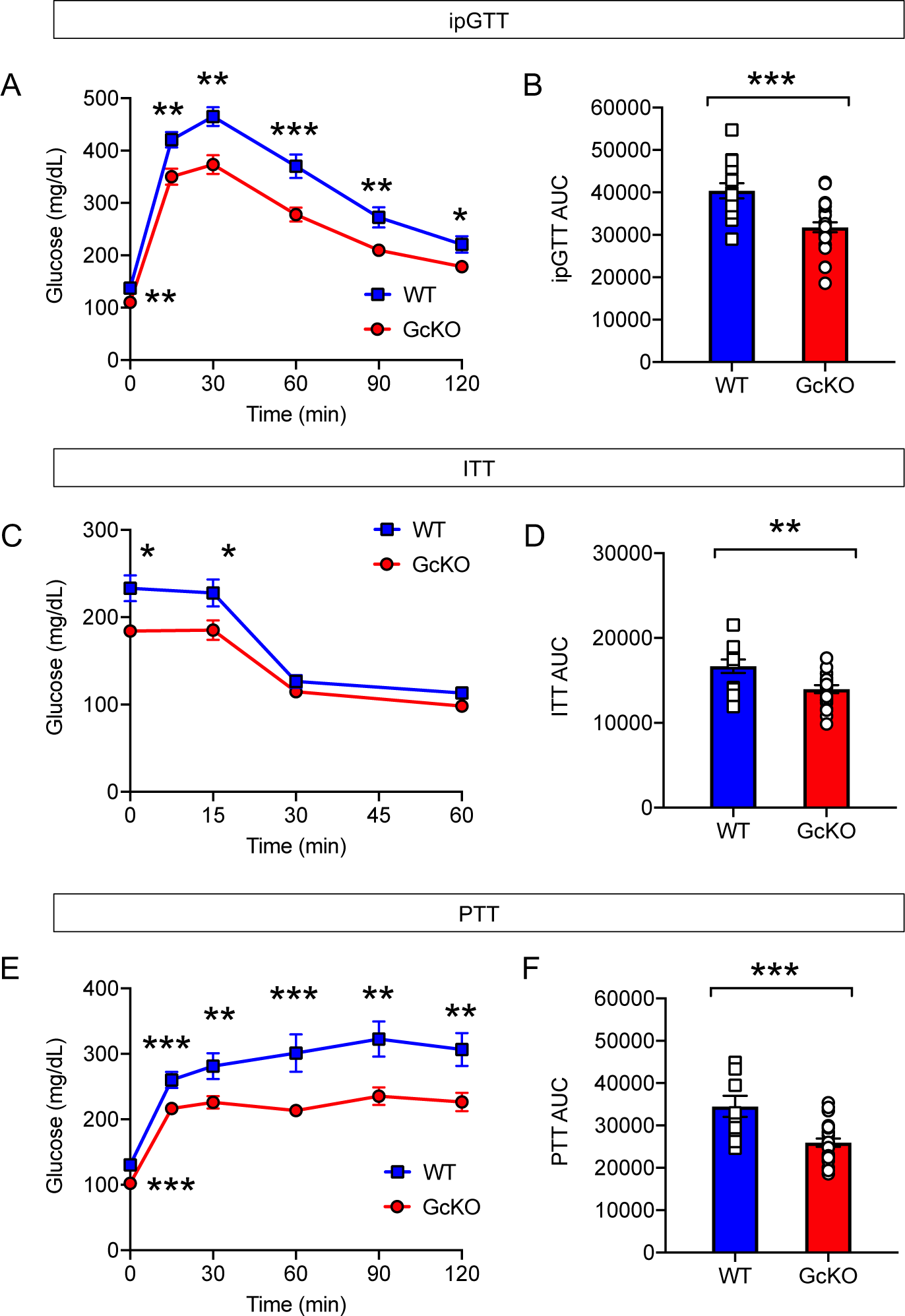
Gc ablation improves HFD-induced glucose and insulin intolerance. **(A)** Intraperitoneal glucose tolerance test performed in HFD-fed WT (n=14) and GcKO (n=25) mice. **(B)** Area under the curve (AUC) in A. **(C)** Insulin tolerance test performed in HFD-fed WT (n=12) and GcKO (n=21) mice. **(D)** AUC in C. **(E)** Pyruvate tolerance test performed in HFD-fed WT (n=10) and GcKO (n=23) mice. **(F)** AUC in E. Error bars represent ±SEM, \**p* <0.05, \*\**p* <0.01, \*\*\**p* <0.005 by Student’s *t* test.

In insulin tolerance tests, we found that the glucose-lowering effects of insulin were similar, although the differences in fasting glucose values between WT vs. GcKO mice render the interpretation of this finding more complex (Fig. 2C-D). In contrast, pyruvate tolerance tests showed a consistently lower glycemic level throughout the 2-hr time frame in GcKO mice, suggesting a significant decrease in the ability to convert pyruvate to glucose in the liver compared to WT (Fig. 2E-F).

### Hyperinsulinemic-euglycemic clamps

To directly assess insulin sensitivity *in vivo*, we performed hyperinsulinemic-euglycemic clamps in HFD-fed WT and GcKO mice (20). We clamped glucose levels at ∼125 mg/dL using a constant insulin infusion at a rate of 2.5 mU/kg/min, and measured the glucose infusion rate necessary to maintain euglycemia (Fig. S3A). We found a threefold higher glucose infusion rate in GcKO mice compared WT (17.6±4.1 *vs*. 5.2±1.9 mg/kg/min, *p* <0.03) (Fig. 3A). Basal hepatic glucose production (HGP) was slightly increased in GcKO mice, and was suppressed by ∼24% following insulin clamp. In contrast, WT mice failed to respond to insulin to any meaningful extent (*p* <0.03) (Fig. 3B). Glucose turnover nearly doubled in GcKO mice compared to WT (29.1±3.6 *vs*. 18.1±1.4 mg/kg/min, *p* <0.03) (Fig. 3C). GcKO mice also showed a trend of increased glycogen synthesis that did not reach statistical significance (Fig. 3D), while glycolysis was similar in mice of both genotypes (data not shown). Overall, these data indicate that Gc ablation is associated with preserved insulin sensitivity following HFD, most notably with regard to insulin’s ability to suppress HGP.

**Figure 3.**
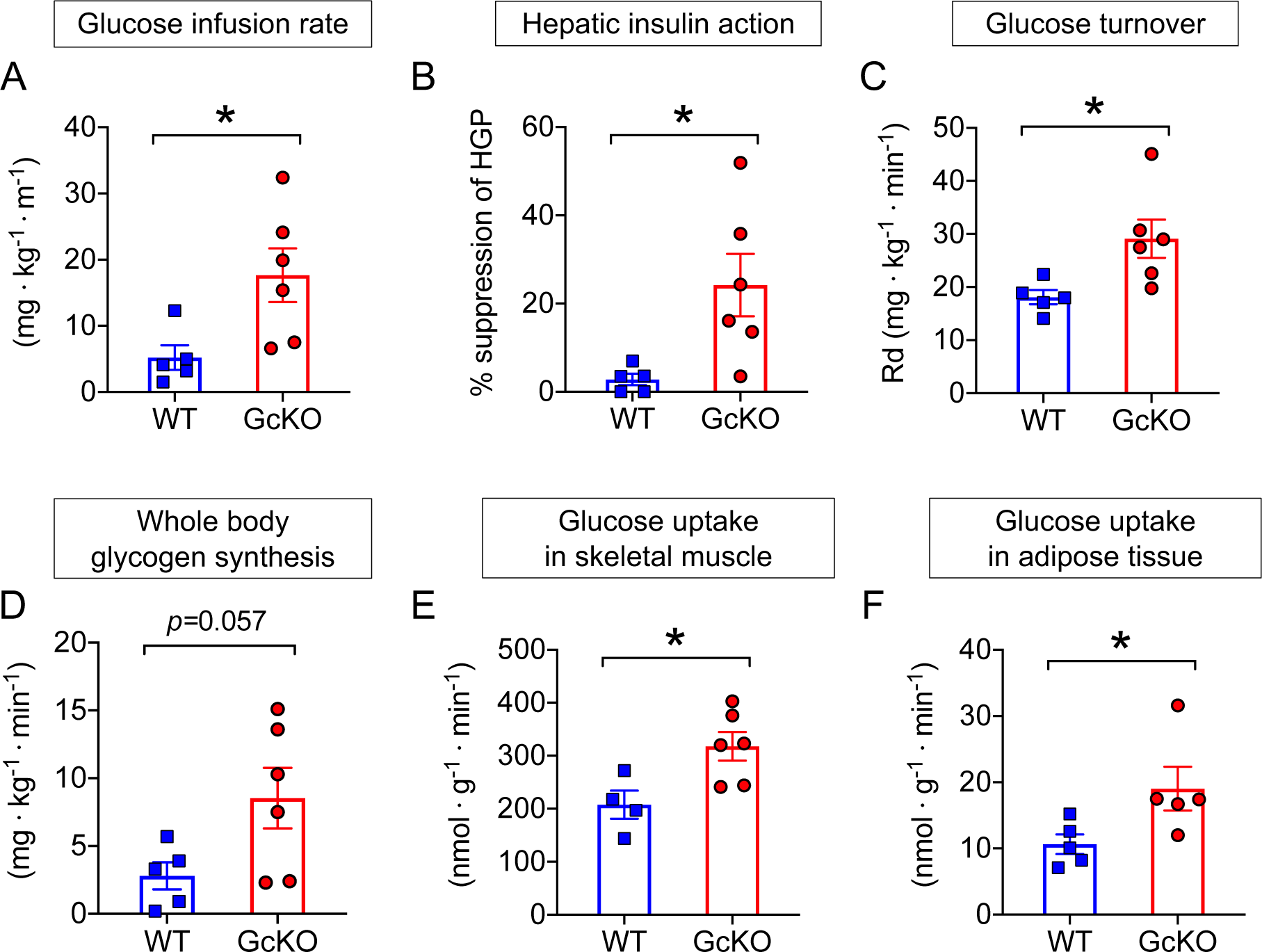
Hyperinsulinemic-euglycemic clamps. **(A)** Glucose infusion rate (GIR) in HFD-fed WT and GcKO mice during clamps. **(B)** Hepatic insulin action, calculated as the ratio of clamp hepatic glucose production (HGP) to basal HGP, and presented as percent suppression. **(C)** Rate of glucose disposal (Rd). **(D)** Whole body glycogen synthesis. **(E)** Glucose uptake in gastrocnemius muscle. **(F)** Glucose uptake in epidydimal fat pad. WT n=5, GcKO n=6. Error bars represent ±SEM, \**p* <0.05 by Student’s *t* test.

### Glucose uptake in muscle and adipose tissue

Skeletal muscle is the major site of insulin-stimulated glucose uptake in humans (22). We examined glucose uptake in skeletal muscle during hyperinsulinemic-euglycemic clamps. We found significantly increased glucose uptake in gastrocnemius of GcKO compared to WT mice (318±27 *vs*. 208±27 nmol/g/min, *p* <0.03) (Fig. 3E). White adipose tissue (WAT) contributes to about 10% of whole-body glucose uptake in response to insulin. Therefore, we also examined glucose uptake in epidydimal WAT, and found a significant increase in GcKO compared to WT (19±3.3 *vs*. 10.6±1.5 nmol/g/min, *p* <0.05) (Fig. 3F). These data demonstrate that in addition to hepatic insulin sensitivity, Gc ablation preserves insulin action on skeletal muscle and subcutaneous adipose tissue glucose uptake. Moreover, we assessed hepatic triglyceride and glycogen content at the end of clamp studies. While there is no difference in triglyceride levels (Fig. S3B), we found ∼50% reduction in hepatic glycogen content in GcKO compared to WT (Sig. S3C). This finding corroborated with increased insulin action in skeletal muscle and adipose tissue.

### Lipid profile in GcKO mice

Increased levels of plasma non-esterified fatty acid (NEFA) impair insulin’s ability to suppress HGP (23), and can cause insulin resistance (24), as well as hepatic lipid accumulation. We found that NEFA levels were decreased by ∼17% in GcKO compared to WT (Fig. 4A). In addition, plasma triglyceride levels are generally elevated in poorly controlled diabetics as well as in pre-diabetes, and are thought to predispose to atherosclerosis (25). TG levels also showed a ∼35% decrease in GcKO mice compared to WT (Fig. 4B). In contrast, we did not find differences in total cholesterol plasma levels (Fig. 4C).

**Figure 4.**
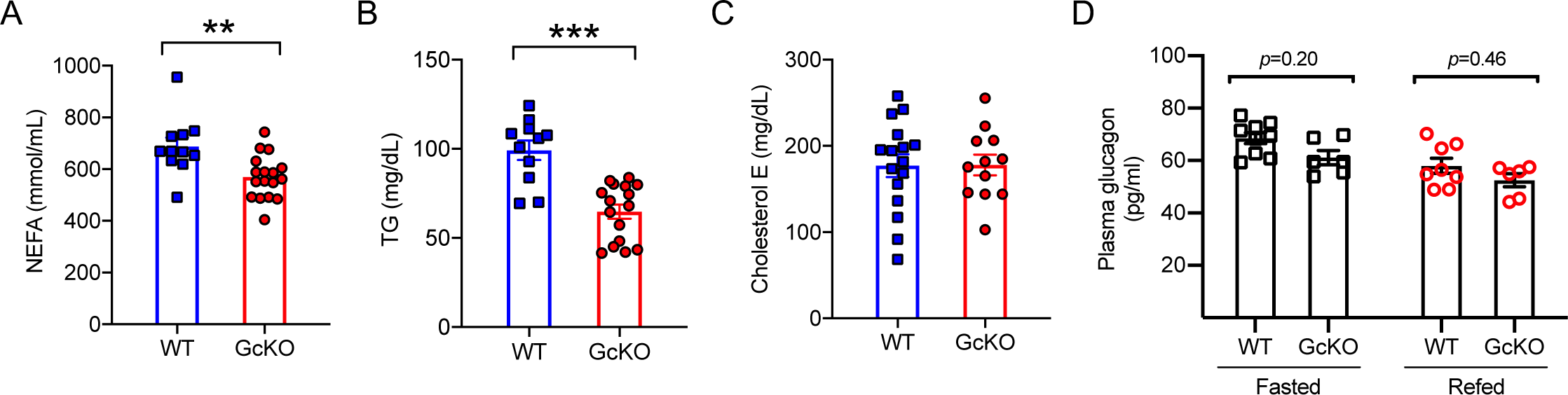
Circulating lipid levels. Plasma non-esterified fatty acid (NEFA) **(A**), triglyceride (TG) **(B)**, and cholesterol E levels **(C)** in HFD-fed WT and GcKO mice. WT n ≥11, GcKO n ≥12. **(D)** Plasma glucagon levels in fasted or refed WT and GcKO mice. WT n= 8, GcKO n= 6. Error bars represent ± SEM, \*\**p* <0.01, \*\*\**p* <0.005 by Student’s *t* test.

To assess if the decreased levels of NEFA and TG are a consequence of decreased circulating glucagon during the fasted state, we measure plasma glucagon levels in WT and GcKO mice. While refeeding lowered glucagon secretion in both WT and GcKO, we did not find differences in fasted or refed state between genotypes (Fig. 4D). These data indicated that Gc ablation did not alter glucagon secretion, and the changes in lipid profiles in GcKO mice are not related to glucagon levels.

### Gene expression survey

To determine whether the preserved insulin sensitivity in GcKO mice was accompanied by changes in gene expression, we measured mRNA levels of several key metabolic genes in liver, the main site of Gc expression (Fig. 5A). RT-qPCR determined that *Gc* mRNA was effectively ablated (Fig. 5B). Levels of mRNA encoding the gluconeogenic rate-limiting enzyme, *Pck1*, were decreased in the fasted state in GcKO mice, while levels of *G6pc* and *Fbp1* were decreased in both fasted and refed conditions compared to WT mice (Fig. 5C-E), consistent with preserved hepatic insulin action found during the euglycemic clamps. In contrast, there was no difference in the glycolytic gene *Gck* (Fig. 5F) or in the glycogen synthase inhibitor, Gsk3β (Fig. 5G).

**Figure 5.**
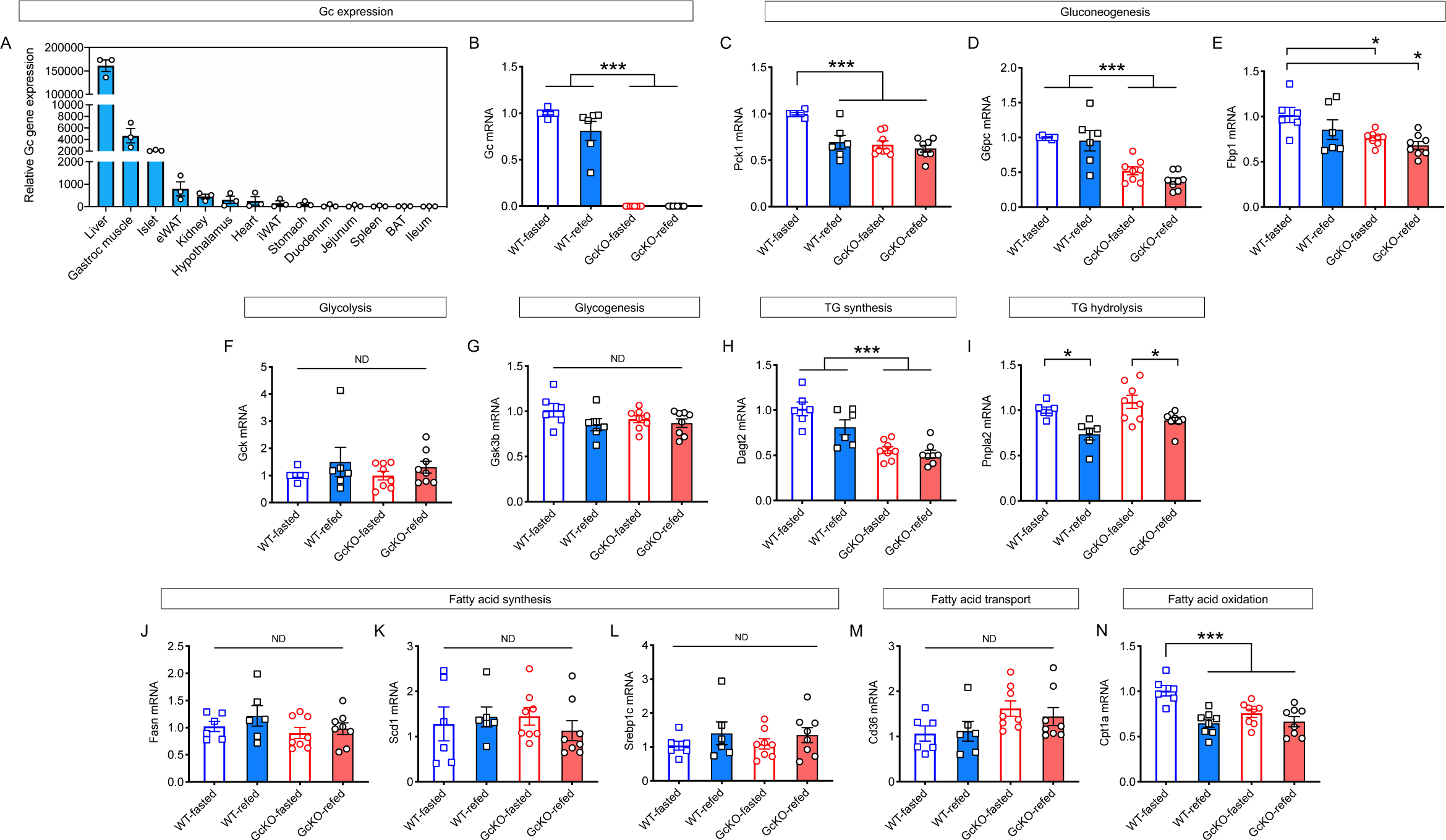
Gene expression analysis. **(A)** Relative *Gc* expression in various metabolic tissues. **(B-N)** Gene expression in fasted or refed WT and GcKO mice: **(B)** Gc; **(C-E)** gluconeogenic genes *Pck1* (C), *G6pc* (D), and *Fbp1* (E); **(F)** *Gck*; **(G)** *Gsk3*β; **(H)** *Dgat2***; (I)** *Pnpla2*; **(J-L)** fatty acid synthesis genes *Fasn* (J), *Scd1* (K), and *Srebp1c* (L); **(M)** fatty acid transporter *Cd36*; and **(N)** *Cpt1a*. WT n=6; GcKO n=8. Error bars represent ± SEM, \**p* <0.05, \*\**p* <0.01, \*\*\**p* <0.005 by Student’s *t* test or ANOVA.

With regard to lipid synthetic/turnover enzymes, we found a reduction of mRNA encoding the TG synthetic enzyme, *Dgat2*, in fasted and refed GcKO mice (Fig. 5H), but no difference in the levels of the TG hydrolytic enzyme mRNA, *Pnpla2* (Fig. 5I), or fatty acid synthesis genes *Fasn, Scd1*, or *Srebp1*, as well as fatty acid transporter, *Cd36* (Fig. 5J-M). We observed a significant reduction in mRNA levels encoding the mitochondrial fatty acid oxidation enzyme *Cpt1a* in fasted and refed GcKO compared to fasted WT mice (Fig. 5N). Thus, changes to gene expression are consistent with the metabolic changes showing GcKO decreased HGP, TG, and NEFA levels, but no changes in glycolysis or glycogen synthesis.

We also measured *Fasn, Scd1, Cpt1a*, and *Pnpla2* mRNA levels in visceral/epidydimal and subcutaneous/inguinal adipose tissue, and found that *Fasn* is decreased in both fat pads, while *Scd1* mRNA is lower in visceral fat pad (Fig. S4A-H). These data are compatible with the possibility that GcKO mice have lower adipose tissue lipogenesis, consistent with the observed reduced fat mass in GcKO mice.

### Acute lowering of Gc levels improves glucose tolerance in HFD-fed mice

To investigate the underlying mechanism of improved insulin sensitivity in HFD-fed GcKO mice, we examined insulin-signaling substrates in the liver. We found increased levels of phosphorylated Akt at serine 473 normalized to total Akt in GcKO mice after refeeding (Fig. 6A, 6B, S5A, and S5B), consistent with augmented hepatic insulin action during hyperinsulinemic-euglycemic clamps (Fig. 3B).

**Figure 6.**
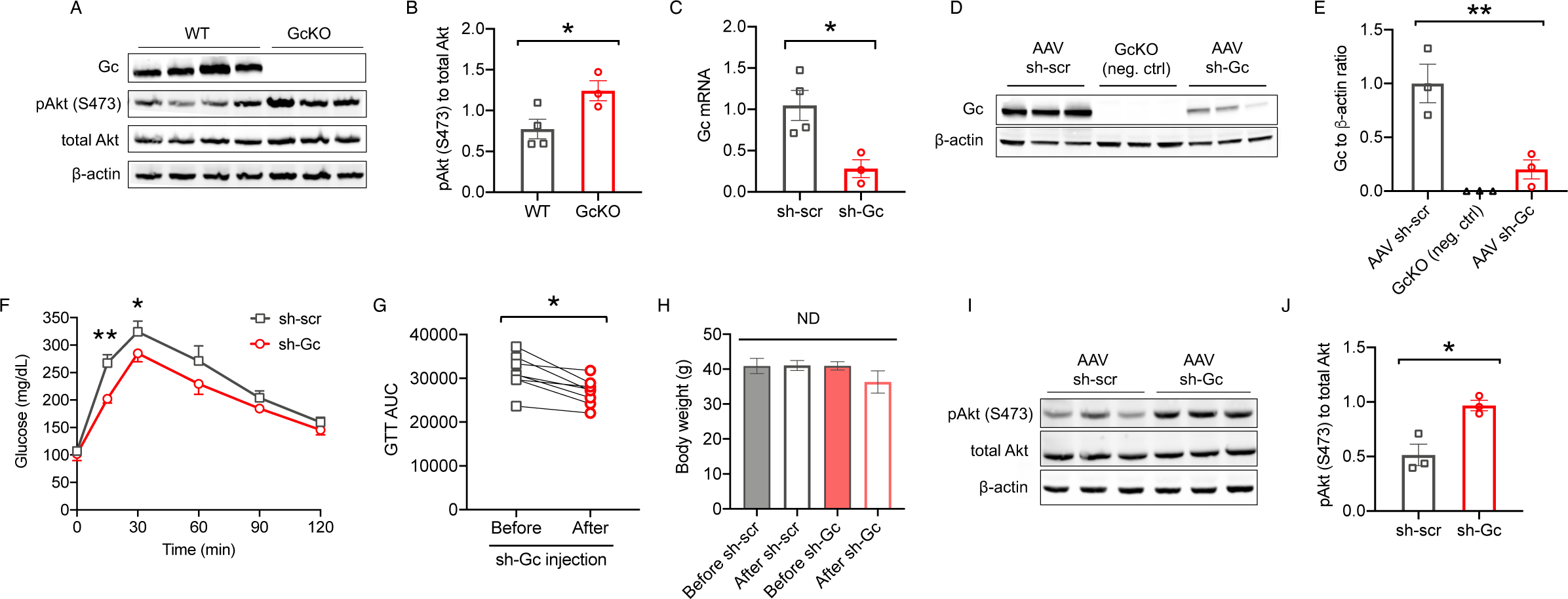
Inhibition of hepatic Gc with adeno-associated virus (AAV). **(A)** Immunoblots showing Gc and insulin signaling substrates in HFD-fed WT and GcKO mice. **(B)** Quantification of (A). **(C)** Gc expression in mice injected with AAV encoding shRNA against control (sh-scr) or *Gc* (sh-Gc). **(D)** Immunoblotting of liver Gc and internal control β-actin in WT mice injected with AAV carrying sh-scr or sh-Gc. GcKO mice were used as a negative control. **(E)** Quantification of (D). **(F)** Glucose tolerance test in WT mice injected with control (sh-scr, n= 4) or Gc (sh-Gc, n= 8) shRNA. **(G)** Area under the curve from glucose tolerance test in individual WT mice before and after adeno-associated viral sh-Gc injection (n= 8). **(H)** Body weight of mice before and after sh-scr (n= 4) or sh-Gc (n= 8) AAV injection. **(I)** Immunoblotting of insulin signaling substrates in the liver of mice receiving sh-scr or sh-Gc AAV injection. **(J)** Quantification of (I). ND indicates no difference. Error bars represent ±SEM, \**p* <0.05, \*\**p* <0.01 by Student’s *t* test.

The changes observed in GcKO mice may reflect adaptive changes occurring in response to the constitutive lack of this protein. To investigate whether an acute inhibition of Gc levels can improve insulin sensitivity, we injected adeno-associated virus (AAV8) carrying shRNA against Gc (sh-Gc) or control (sh-scr) to inhibit Gc production in the liver of HFD-fed mice. Western blotting analyses confirmed that sh-Gc decreased Gc mRNA (Fig. 6C) and protein levels (Fig. 6D and 6E) by ∼70%. Three weeks post-injection, we performed glucose tolerance test, and found a significant improvement in glucose tolerance in sh-Gc mice compared to sh-scr (Fig. 6F). Analyses of individual mice before and after AAV injection indicated that the decrease occurred across the board in virtually all mice examined (Fig. 6G). In contrast, no significant difference was observed in sh-scr injected mice (Fig. S5C). Moreover, this improvement of glucose tolerance was not accompanied by weight loss (Fig. 6H). This result suggests that, the beneficial effects seen in *Gc* null mice have body weight-dependent and body weight-independent components. We observed elevated hepatic phosphorylated-Akt levels in mice receiving sh-Gc injection (Fig. 6I and 6J). Interestingly, acute ablation of Gc improved plasma insulin levels after refeeding (Fig. S5D), while maintaining similar lipid profile (Fig. S5E-G). These data suggest the possibility of treatments for insulin type 2 diabetes based on inhibition of Gc levels.

## DISCUSSION

The main findings of this work are: *i*) Gc ablation in mice reduced fat mass and body weight following HFD, and lowered plasma NEFA and TG levels; *ii*) Gc-deficient mice also showed improved insulin sensitivity in liver, adipose tissue, and skeletal muscle; *iii*) acute inhibition of Gc, achieved using adeno-associated virus to target hepatic Gc mRNA, improved glucose tolerance in HFD-fed mice. We previously reported that Gc is a pancreatic α-cell signature gene, expression of which is activated in dedifferentiating, dysfunctional β-cells during the progression of multiparity-associated diabetes (14). Ablation of Gc resulted in increased insulin secretion in hyperglycemic clamps in HFD-fed animals and in isolated islets, consistent with a paracrine/autocrine role of Gc to contribute to diabetic β-cell failure. The present data add a new and unforeseen dimension to the gamut of Gc functions by demonstrating that its ablation preserves insulin sensitivity when mice are fed a metabolically unhealthy diet. Thus, Gc inhibition has a dual effect to maintain β-cell insulin secretion and insulin sensitivity, achieving a target profile that has long been viewed as ideal for novel therapeutics in type 2 diabetes. Indeed, none of the currently approved anti-diabetic drugs achieves both goals in one fell swoop. Adding to the potential translational interest of these observations is the fact that Gc can be targeted either as a circulating factor, or through hepatocyte-directed RNA-based therapeutics.

Gc is primarily a vitamin D binding protein (12). ∼85% of circulating 25-hydroxyvitamin D [25(OH)D] is bound to Gc, with the remainder weakly bound to albumin, from which it becomes dissociated in order to enter tissues. Only ∼0.03% of 25(OH)D circulates in blood as free form and has direct access to target cells. Both the free and albumin-bound 25(OH)D fractions are viewed as bioavailable and engage in their metabolic function (26). In contrast, Gc-bound vitamin D and its metabolites are biologically unavailable, with the exception of tissues such as the kidney that express the megalin/cubilin transport system, which allows for Gc-bound 25(OH)D to enter cells (27). The half-life of circulating Gc is 2.5 to 3 days. Gc possesses a single binding site for all vitamin D metabolites, including active form of vitamin D-1,25 (OH)_2_D and the parental vitamin D, with highest affinity for the major form of plasma vitamin D-25(OH)D (12). Although Gc-bound 25(OH)D is biologically unavailable, Gc acts as a reservoir for circulating 25(OH)D by increasing their half-life, as 25(OH)D was rapidly metabolized and excreted in the urine in the absence of Gc (28). While total concentrations of 25(OH)D and 1,25(OH)_2_D in Gc-deficient mice (28) or in a human nullizygous *GC* mutant (29) are extremely low, calcium and bone homeostasis remain relatively normal. As such, only when placed on a vitamin D deficient diet for four weeks that *Gc* null mice showed a mild bone mineralization defect (28). Furthermore, Gc plays a role in transporting fatty acids (30). These findings are consistent with the possibility that there are other functions of Gc, in addition to transporting vitamin D.

The relationship between diabetes and vitamin D levels has long been debated. Several observational and physiological studies have linked vitamin D insufficiency with β-cell dysfunction, insulin resistance, and obesity, both in adults and children (31). However, the pivotal D2d trial, a randomized control trial in individuals at risk of developing diabetes, showed no statistically significant benefit from vitamin D supplementation (32), as did similar trials (33,34). Thus, the balance of probabilities is that the effect of Gc deficiency is independent of vitamin D levels, especially since the latter tend to be low in the plasma when Gc is ablated.

A report of a 58-year-old woman born to consanguineous parents with a homozygous null mutation of *GC* confirmed its role as a vitamin D carrier (29). The patient had a history of musculoskeletal pain and fragility fractures, associated with extremely low plasma levels of vitamin D and its metabolites. Although she had diabetes, it was well controlled on metformin (J. Marcadier, personal communication), and was associated with additional comorbidities, including hypertension, dyslipidemia, nonalcoholic fatty liver disease, and autoimmunity, that may have metabolic effects of their own. It should be emphasized that our experiments show beneficial effects of ∼70% Gc inhibition in mice, suggesting that the effective range of inhibition necessary for metabolic benefits can be safe in humans.

We are left with the question of how Gc deficiency promotes insulin sensitivity. There are precedents for circulating proteins that modulate insulin action, for example adiponectin (35), Resistin (36), and Rbp4 (37). The beneficial effects of Gc ablation in HFD-fed GcKO mice run the gamut from decreased weight and fat mass to decreased HGP, NEFA, TG, and increased insulin action on glucose transport. This combination of actions is most consistent with a systemic insulin-like property associated with Gc ablation. One possibility is that Gc carries one or more biochemical entities that confer insulin resistance, and thus its ablation confers protection against this agent(s). Alternatively, Gc can act to prop up insulin receptor signaling, either by acting at the cell membrane or through some heretofore unrecognized intracellular mechanism. However, the latter would require that Gc be taken up by cells, and this property appears to be limited to the kidney. More work will be required to address this question.

From a therapeutic standpoint, there remains an unmet need for insulin sensitizers (38). TZDs, whose use has fallen out of fashion due to a combination of adverse side effects and regulatory setbacks (39), remain the only class of drugs consistently associated with an effect on diabetes prevention (9-11). They also show consistently lower secondary failure rates than metformin, sulfonylureas (40), and Glp1-RA (41). In addition to insulin sensitization, Gc inhibition appears to also protect against β-cell dysfunction and increase insulin secretion, at least in mice. There is a need for therapeutics that can address both β-cell function and peripheral insulin action, and we propose Gc as a candidate target.

## ACKNOWLEDGMENTS

We thank members of the Accili laboratory for discussions, and Thomas Kolar and Ana Flete-Castro (Columbia University) for technical support. We also thank Drs. Jason K. Kim and Randall Friedline (National MMPC at UMass) for clamp studies. Dr. Taiyi Kuo is the guarantor of this work and, as such, had full access to all the data in the study and takes responsibility for the integrity of the data and the accuracy of the data analysis.

## AUTHOR CONTRIBUTIONS

T.K. designed the studies, performed experiments, analyzed the data, and wrote the manuscript. D.A. designed the studies, analyzed the data, and wrote the manuscript.

## SUPPLEMENTAL FIGURE LEGENDS

**Figure S1.**
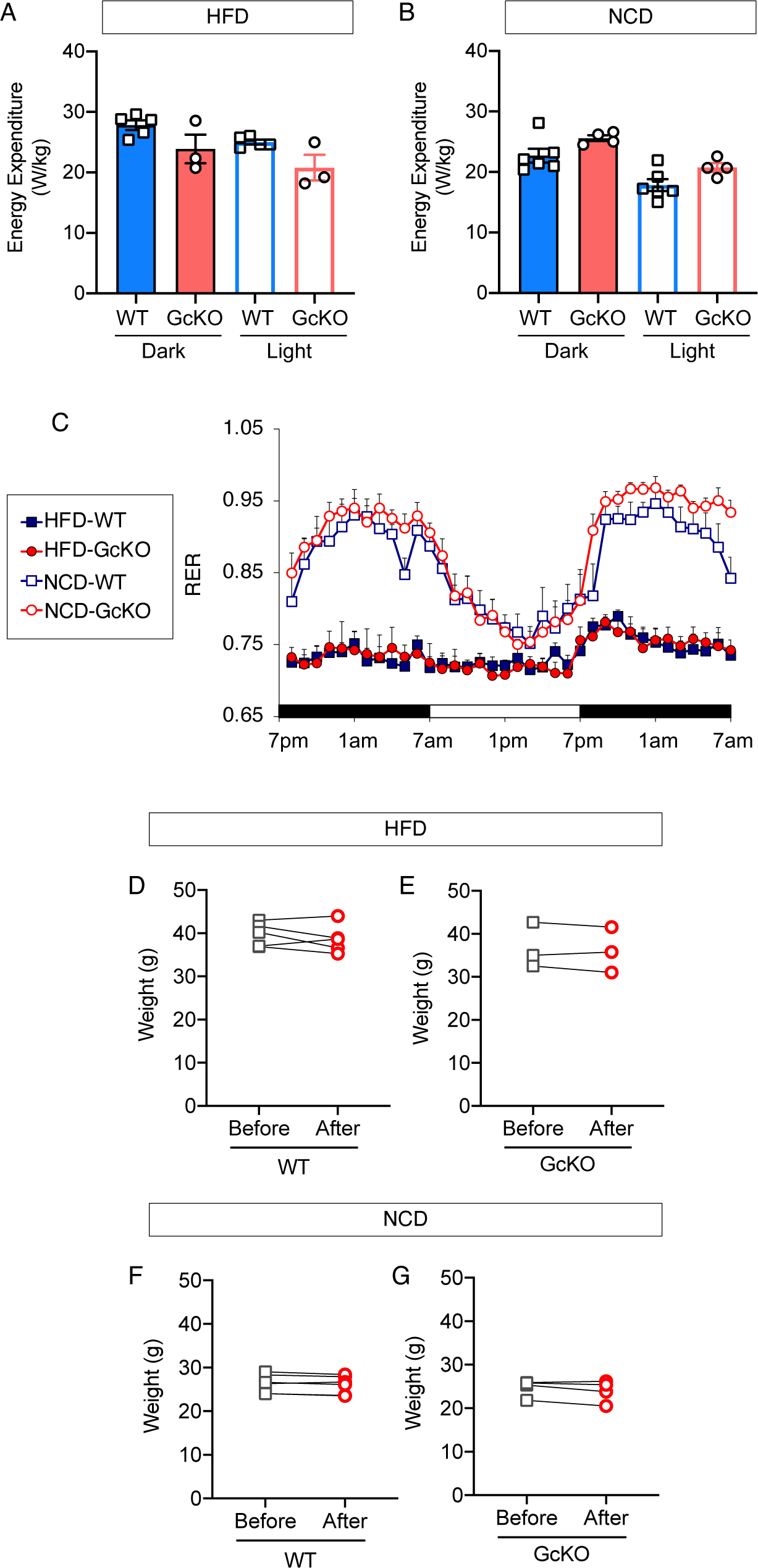
Indirect calorimetry in WT and GcKO mice. Energy expenditure in **(A)** high fat diet (HFD)-fed, or **(B)** normal chow diet (NCD)-fed mice. **(C)** Respiratory exchange rate (RER) in HFD- or NCD-fed WT and GcKO mice. **(D-G)** Body weight before and after indirect calorimetry in (D) HFD-fed WT, (E) HFD-fed GcKO, (F) NCD-fed WT, and (G) NCD-fed GcKO mice.

**Figure S2.**
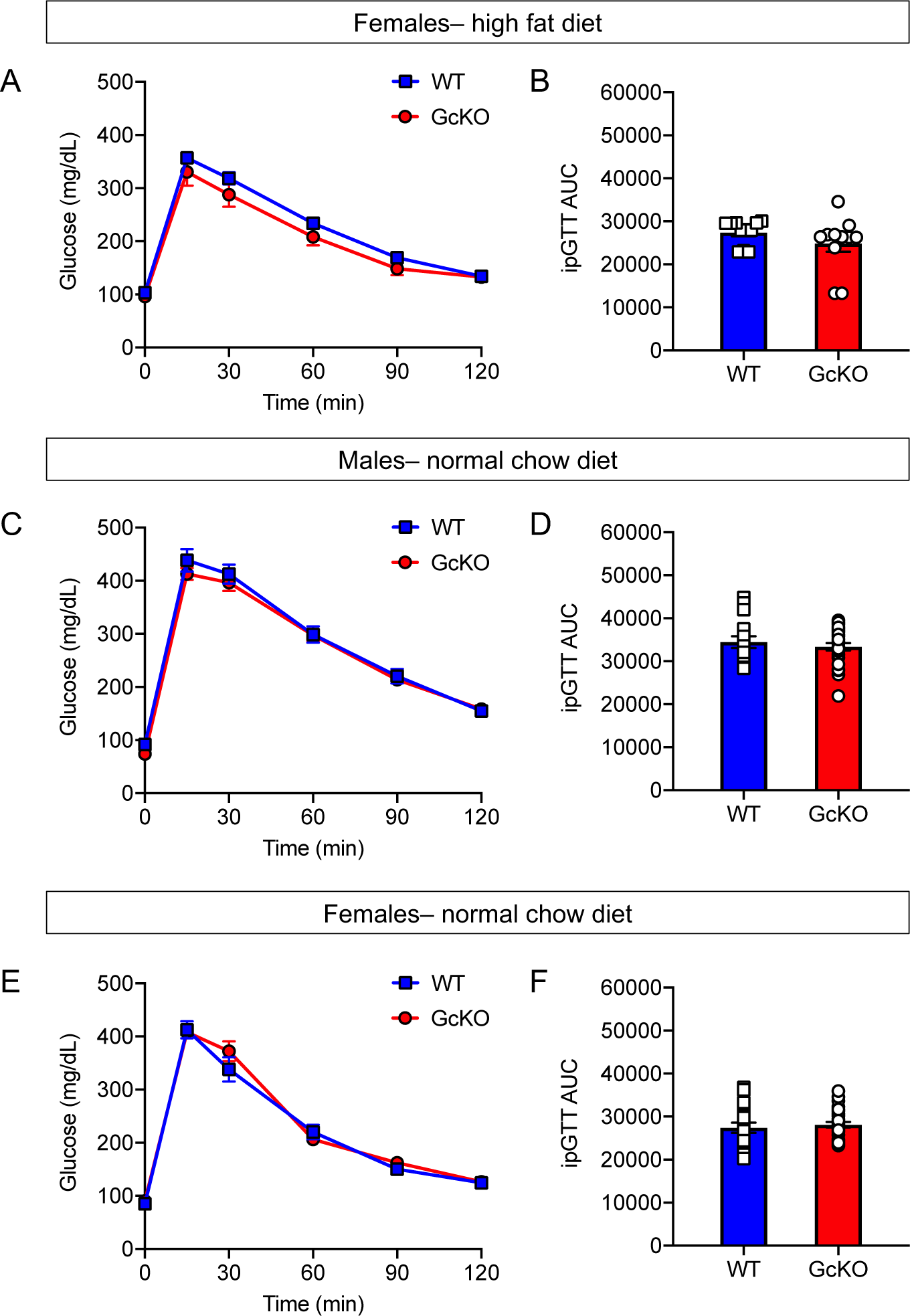
Metabolic characteristics of WT and GcKO mice. **(A)** Intraperitoneal glucose tolerance test (ipGTT) in HFD-fed female WT (n=11) and GcKO (n=11) mice. **(B)** Area under the curve (AUC) in A. **(C)** ipGTT in NCD-fed male WT (n=15) and GcKO (n=25) mice. **(D)** AUC in C. **(E)** ipGTT in NCD-fed female WT (n=18) and GcKO (n=31) mice. **(F)** AUC in E. Error bars represent ±SEM.

**Figure S3.**
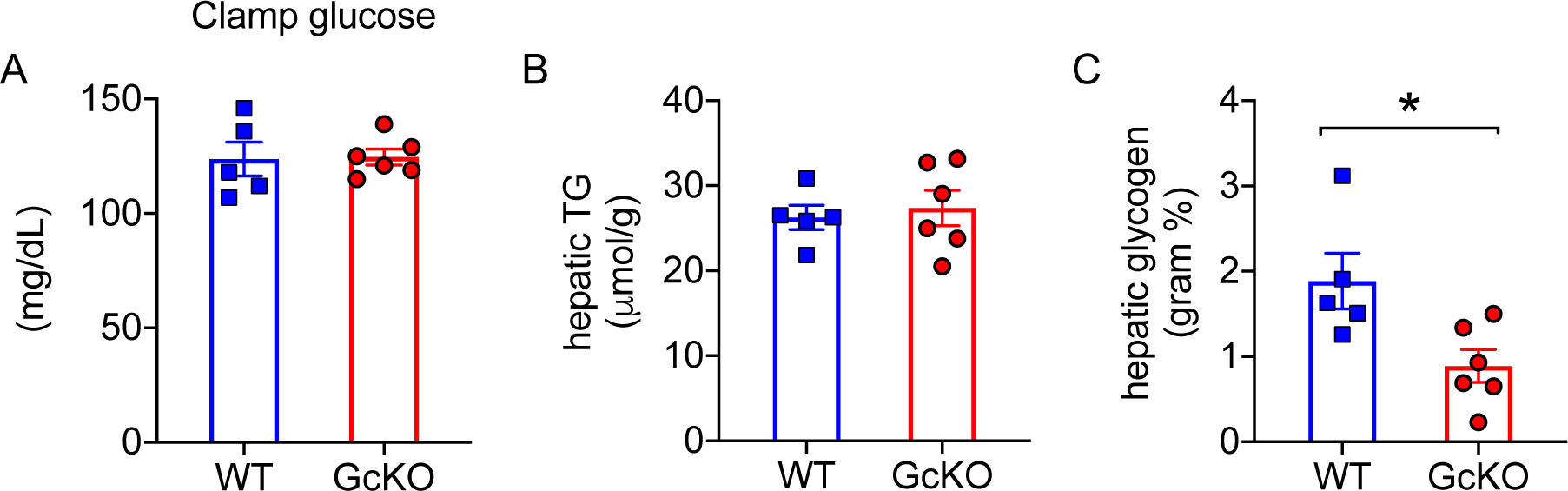
Hepatic TG and glycogen contents. **(A)** Glucose levels during hyperinsulinemic-euglycemic clamps in HFD-fed WT and GcKO mice. **(B)** Post-clamp hepatic TG content. **(C)** Post-clamp hepatic glycogen content. WT n= 5, GcKO n= 6. Error bars represent ±SEM, \**p* <0.05 by Student’s *t* test.

**Figure S4.**
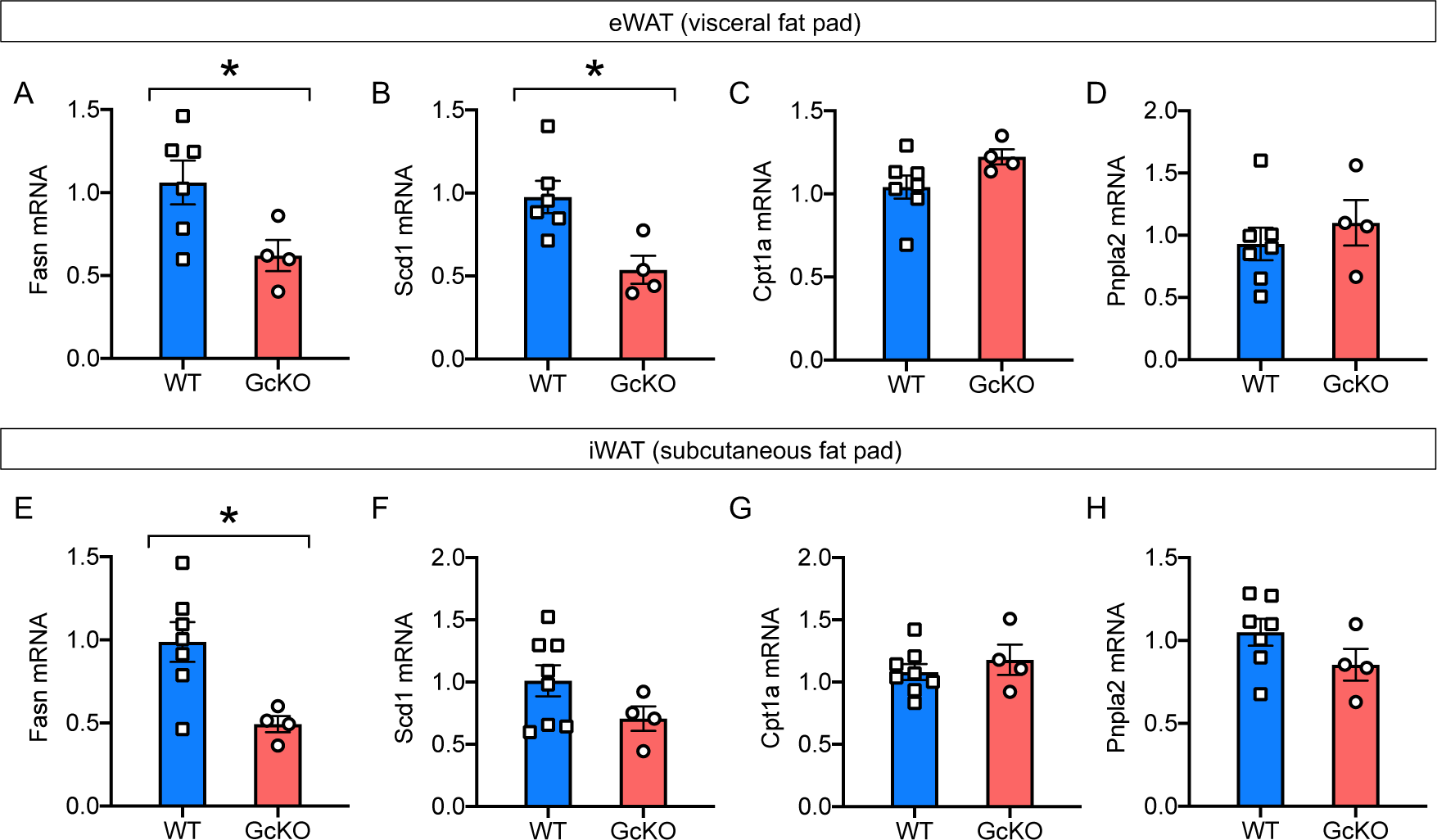
Gene expression analysis in white adipose tissue (WAT). **(A-D)** In epidydimal WAT, a visceral fat pad, mRNA expression of (A) *Fasn*, (B) *Scd1*, (C) *Cpt1a*, and (D) *Pnpla2*. **(E-H)** in inguinal WAT, a subcutaneous fat pad, mRNA expression of (E) *Fasn*, (F) *Scd1*, (G) *Cpt1a*, and (H) *Pnpla2*. WT n= 6, GcKO n= 4. Error bars represent ±SEM, \**p* <0.05 by Student’s *t* test.

**Figure S5.**
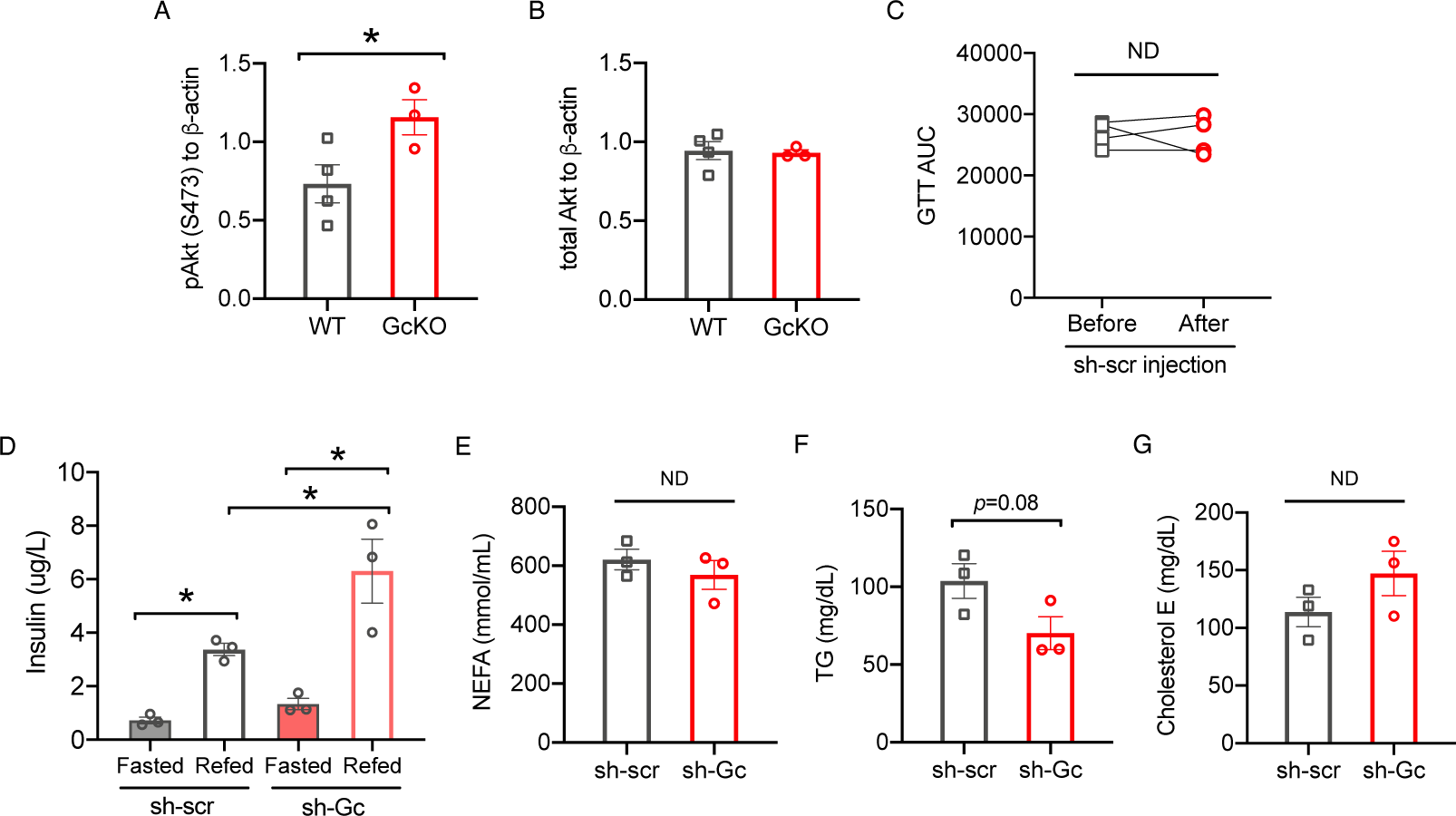
Acute inhibition of hepatic Gc. **(A)** Quantification of normalized phosphorylated Akt at serine 473 to β-actin in HFD-fed WT or GcKO mice. **(B)** Quantification of normalized total Akt to β-actin in HFD-fed WT or GcKO mice. **(C)** Area under the curve from glucose tolerance test in individual WT mice before and after adeno-associated viral sh-scr (n= 4) injection. **(D)** Plasma insulin levels in sh-scr and sh-Gc mice in fasted and refed conditions. **(E-G)** Circulating levels of (E) non-esterified fatty acid (NEFA), (F) triglyceride, and (G) cholesterol E in fasted sh-scr and sh-Gc mice. Error bars represent ±SEM, \**p* <0.05 by Student’s *t* test.

## REFERENCES

1. Accili, D. (2018) Insulin Action Research and the Future of Diabetes Treatment: The 2017 Banting Medal for Scientific Achievement Lecture. Diabetes 67, 1701–1709

2. Weyer, C., Bogardus, C., Mott, D. M., and Pratley, R. E. (1999) The natural history of insulin secretory dysfunction and insulin resistance in the pathogenesis of type 2 diabetes mellitus. J Clin Invest 104, 787–794

3. Lillioja, S., Mott, D. M., Spraul, M., Ferraro, R., Foley, J. E., Ravussin, E., Knowler, W. C., Bennett, P. H., and Bogardus, C. (1993) Insulin resistance and insulin secretory dysfunction as precursors of non-insulin-dependent diabetes mellitus. Prospective studies of Pima Indians. N Engl J Med 329, 1988–1992

4. Brunzell, J. D., Robertson, R. P., Lerner, R. L., Hazzard, W. R., Ensinck, J. W., Bierman, E. L., and Porte, D., Jr. (1976) Relationships between fasting plasma glucose levels and insulin secretion during intravenous glucose tolerance tests. J Clin Endocrinol Metab 42, 222–229

5. Ferrannini, E. (2010) The stunned beta cell: a brief history. Cell metabolism 11, 349–352

6. American Diabetes, A. (2020) 6. Glycemic Targets: Standards of Medical Care in Diabetes-2020. Diabetes Care 43, S66–S76

7. Wilkinson, S., Douglas, I., Stirnadel-Farrant, H., Fogarty, D., Pokrajac, A., Smeeth, L., and Tomlinson, L. (2018) Changing use of antidiabetic drugs in the UK: trends in prescribing 2000-2017. BMJ Open 8, e022768

8. Holst, J. J. (2019) From the Incretin Concept and the Discovery of GLP-1 to Today’s Diabetes Therapy. Front Endocrinol (Lausanne) 10, 260

9. Buchanan, T. A., Xiang, A. H., Peters, R. K., Kjos, S. L., Berkowitz, K., Marroquin, A., Goico, J., Ochoa, C., and Azen, S. P. (2000) Response of pancreatic beta-cells to improved insulin sensitivity in women at high risk for type 2 diabetes. Diabetes 49, 782–788

10. Defronzo, R. A., Tripathy, D., Schwenke, D. C., Banerji, M., Bray, G. A., Buchanan, T. A., Clement, S. C., Gastaldelli, A., Henry, R. R., Kitabchi, A. E., Mudaliar, S., Ratner, R. E., Stentz, F. B., Musi, N., Reaven, P. D., and Study, A. N. (2013) Prevention of diabetes with pioglitazone in ACT NOW: physiologic correlates. Diabetes 62, 3920–3926

11. Kernan, W. N., Viscoli, C. M., Furie, K. L., Young, L. H., Inzucchi, S. E., Gorman, M., Guarino, P. D., Lovejoy, A. M., Peduzzi, P. N., Conwit, R., Brass, L. M., Schwartz, G. G., Adams, H. P., Jr., Berger, L., Carolei, A., Clark, W., Coull, B., Ford, G. A., Kleindorfer, D., O’Leary, J. R., Parsons, M. W., Ringleb, P., Sen, S., Spence, J. D., Tanne, D., Wang, D., Winder, T. R., and Investigators, I. T. (2016) Pioglitazone after Ischemic Stroke or Transient Ischemic Attack. N Engl J Med 374, 1321–1331

12. Bouillon, R., Van Baelen, H., Rombauts, W., and De Moor, P. (1976) The purification and characterisation of the human-serum binding protein for the 25-hydroxycholecalciferol (transcalciferin). Identity with group-specific component. Eur J Biochem 66, 285–291

13. Wang, Y. J., Schug, J., Won, K. J., Liu, C., Naji, A., Avrahami, D., Golson, M. L., and Kaestner, K. H. (2016) Single cell transcriptomics of the human endocrine pancreas. Diabetes

14. Kuo, T., Damle, M., Gonzalez, B. J., Egli, D., Lazar, M. A., and Accili, D. (2019) Induction of alpha cell-restricted Gc in dedifferentiating beta cells contributes to stress-induced beta-cell dysfunction. JCI Insight 5

15. Swisa, A., Avrahami, D., Eden, N., Zhang, J., Feleke, E., Dahan, T., Cohen-Tayar, Y., Stolovich-Rain, M., Kaestner, K. H., Glaser, B., Ashery-Padan, R., and Dor, Y. (2017) PAX6 maintains beta cell identity by repressing genes of alternative islet cell types. J Clin Invest 127, 230–243

16. Ediger, B. N., Du, A., Liu, J., Hunter, C. S., Walp, E. R., Schug, J., Kaestner, K. H., Stein, R., Stoffers, D. A., and May, C. L. (2014) Islet-1 Is essential for pancreatic beta-cell function. Diabetes 63, 4206–4217

17. Kuo, T., Kim-Muller, J. Y., McGraw, T. E., and Accili, D. (2016) Altered Plasma Profile of Antioxidant Proteins as an Early Correlate of Pancreatic beta Cell Dysfunction. J Biol Chem 291, 9648–9656

18. Kuo, T., Chen, T. C., Yan, S., Foo, F., Ching, C., McQueen, A., and Wang, J. C. (2014) Repression of glucocorticoid-stimulated angiopoietin-like 4 gene transcription by insulin. J Lipid Res 55, 919–928

19. Kuo, T., Kraakman, M. J., Damle, M., Gill, R., Lazar, M. A., and Accili, D. (2019) Identification of C2CD4A as a human diabetes susceptibility gene with a role in beta cell insulin secretion. Proc Natl Acad Sci U S A 116, 20033–20042

20. Kim, J. K. (2009) Hyperinsulinemic-euglycemic clamp to assess insulin sensitivity in vivo. Methods Mol Biol 560, 221–238

21. Zhu, C., Kim, K., Wang, X., Bartolome, A., Salomao, M., Dongiovanni, P., Meroni, M., Graham, M. J., Yates, K. P., Diehl, A. M., Schwabe, R. F., Tabas, I., Valenti, L., Lavine, J. E., and Pajvani, U. B. (2018) Hepatocyte Notch activation induces liver fibrosis in nonalcoholic steatohepatitis. Sci Transl Med 10

22. DeFronzo, R. A., and Tripathy, D. (2009) Skeletal muscle insulin resistance is the primary defect in type 2 diabetes. Diabetes Care 32 Suppl 2, S157–163

23. Roden, M., Stingl, H., Chandramouli, V., Schumann, W. C., Hofer, A., Landau, B. R., Nowotny, P., Waldhausl, W., and Shulman, G. I. (2000) Effects of free fatty acid elevation on postabsorptive endogenous glucose production and gluconeogenesis in humans. Diabetes 49, 701–707

24. Shi, H., Kokoeva, M. V., Inouye, K., Tzameli, I., Yin, H., and Flier, J. S. (2006) TLR4 links innate immunity and fatty acid-induced insulin resistance. J Clin Invest 116, 3015–3025

25. Chait, A., and Goldberg, I. (2017) Treatment of Dyslipidemia in Diabetes: Recent Advances and Remaining Questions. Current diabetes reports 17, 112

26. Chun, R. F., Peercy, B. E., Orwoll, E. S., Nielson, C. M., Adams, J. S., and Hewison, M. (2014) Vitamin D and DBP: the free hormone hypothesis revisited. J Steroid Biochem Mol Biol 144 Pt A, 132–137

27. Nykjaer, A., Dragun, D., Walther, D., Vorum, H., Jacobsen, C., Herz, J., Melsen, F., Christensen, E. I., and Willnow, T. E. (1999) An endocytic pathway essential for renal uptake and activation of the steroid 25-(OH) vitamin D3. Cell 96, 507–515

28. Safadi, F. F., Thornton, P., Magiera, H., Hollis, B. W., Gentile, M., Haddad, J. G., Liebhaber, S. A., and Cooke, N. E. (1999) Osteopathy and resistance to vitamin D toxicity in mice null for vitamin D binding protein. J Clin Invest 103, 239–251

29. Henderson, C. M., Fink, S. L., Bassyouni, H., Argiropoulos, B., Brown, L., Laha, T. J., Jackson, K. J., Lewkonia, R., Ferreira, P., Hoofnagle, A. N., and Marcadier, J. L. (2019) Vitamin D-Binding Protein Deficiency and Homozygous Deletion of the GC Gene. N Engl J Med 380, 1150–1157

30. Williams, M. H., Van Alstyne, E. L., and Galbraith, R. M. (1988) Evidence of a novel association of unsaturated fatty acids with Gc (vitamin D-binding protein). Biochem Biophys Res Commun 153, 1019–1024

31. Wexler, D. J. (2019) D2d - No Defense against Diabetes. N Engl J Med 381, 581–582

32. Pittas, A. G., Dawson-Hughes, B., Sheehan, P., Ware, J. H., Knowler, W. C., Aroda, V. R., Brodsky, I., Ceglia, L., Chadha, C., Chatterjee, R., Desouza, C., Dolor, R., Foreyt, J., Fuss, P., Ghazi, A., Hsia, D. S., Johnson, K. C., Kashyap, S. R., Kim, S., LeBlanc, E. S., Lewis, M. R., Liao, E., Neff, L. M., Nelson, J., O’Neil, P., Park, J., Peters, A., Phillips, L. S., Pratley, R., Raskin, P., Rasouli, N., Robbins, D., Rosen, C., Vickery, E. M., Staten, M., and Group, D. d. R. (2019) Vitamin D Supplementation and Prevention of Type 2 Diabetes. N Engl J Med 381, 520–530

33. Jorde, R., Sollid, S. T., Svartberg, J., Schirmer, H., Joakimsen, R. M., Njolstad, I., Fuskevag, O. M., Figenschau, Y., and Hutchinson, M. Y. (2016) Vitamin D 20,000 IU per Week for Five Years Does Not Prevent Progression From Prediabetes to Diabetes. J Clin Endocrinol Metab 101, 1647–1655

34. Kawahara, T., Suzuki, G., Inazu, T., Mizuno, S., Kasagi, F., Okada, Y., and Tanaka, Y. (2016) Rationale and design of Diabetes Prevention with active Vitamin D (DPVD): a randomised, double-blind, placebo-controlled study. BMJ Open 6, e011183

35. Okada-Iwabu, M., Yamauchi, T., Iwabu, M., Honma, T., Hamagami, K., Matsuda, K., Yamaguchi, M., Tanabe, H., Kimura-Someya, T., Shirouzu, M., Ogata, H., Tokuyama, K., Ueki, K., Nagano, T., Tanaka, A., Yokoyama, S., and Kadowaki, T. (2013) A small-molecule AdipoR agonist for type 2 diabetes and short life in obesity. Nature 503, 493–499

36. Schwartz, D. R., and Lazar, M. A. (2011) Human resistin: found in translation from mouse to man. Trends Endocrinol Metab 22, 259–265

37. Zemany, L., Bhanot, S., Peroni, O. D., Murray, S. F., Moraes-Vieira, P. M., Castoldi, A., Manchem, P., Guo, S., Monia, B. P., and Kahn, B. B. (2015) Transthyretin Antisense Oligonucleotides Lower Circulating RBP4 Levels and Improve Insulin Sensitivity in Obese Mice. Diabetes 64, 1603–1614

38. Kim-Muller, J. Y., and Accili, D. (2011) Cell biology. Selective insulin sensitizers. Science 331, 1529–1531

39. Rosen, C. J. (2010) Revisiting the rosiglitazone story--lessons learned. The New England journal of medicine 363, 803–806

40. Kahn, S. E., Haffner, S. M., Heise, M. A., Herman, W. H., Holman, R. R., Jones, N. P., Kravitz, B. G., Lachin, J. M., O’Neill, M. C., Zinman, B., Viberti, G., and Group, A. S. (2006) Glycemic durability of rosiglitazone, metformin, or glyburide monotherapy. N Engl J Med 355, 2427–2443

41. Home, P. D., Ahren, B., Reusch, J. E. B., Rendell, M., Weissman, P. N., Cirkel, D. T., Miller, D., Ambery, P., Carr, M. C., and Nauck, M. A. (2017) Three-year data from 5 HARMONY phase 3 clinical trials of albiglutide in type 2 diabetes mellitus: Long-term efficacy with or without rescue therapy. Diabetes Res Clin Pract 131, 49–60

